# Novel class of yellow emitting carbon dots stimulate collective cell migration and 3D uptake *in vivo*

**DOI:** 10.1101/2022.07.04.498723

**Authors:** Udisha Singh, Krupa Shah, Krupa Kansara, Ashutosh Kumar, Dhiraj Bhatia

## Abstract

We present a new class of nitrogen-doped yellow fluorescent carbon dots, synthesized using a one-step hydrothermal method. These bright fluorescent nanoparticles have excitation and emission spectra near the red region of the visible light spectrum that are quite useful for bioimaging applications. Using organic molecules like ortho- phenylenediamine (OPDA), L-ascorbic acid and urea, yellow fluorescent carbon dots (CDs) were synthesized. We obtained a scalable number of CDs having an average size of 3 nm. The CDs show significant emission spectra in the yellow fluorescence region (λ_em_= 557 nm). The CDs show remarkable stability in their fluorescence in different pH conditions, ionic stability, photostability as well as thermal stability. These CDs are efficiently uptaken by mammalian cells through clathrin-mediated pathway. Apart from *in vitro* studies we have also used zebrafish larvae as a 3D *in vivo* model, and showed that CDs were uptaken efficiently by larvae showing maximum accumulation and fluorescence in the yolk sac region and the notochord region. The CDs also offer enhancement in cell proliferation, hence showing the application in wound healing. The fluorescence of CDs is quite robust and is not affected by most external stimuli, hence can be explored as a promising bioimaging tool for targeted bioimaging and biomedical applications.

## 1. Introduction

Fullerenes, carbon nanotubes, graphene, graphene subordinates, nanodiamonds, and especially carbon-based quantum dots are such carbon-based nanomaterials that have drawn broad consideration in different disciplines because of their novel underlying chemical and physical properties.^1,2^ Scientists have realized that nanodiamonds are hard to get synthesized and separate; other nanomaterials like carbon nanotubes (CNTs), graphene and fullerenes show inferior water dispersibility and trouble giving optimal fluorescence when excited. These undesirable characteristics have restricted their applications in different working areas.^2^ Nanomaterials, which are non-carbon-based such as semiconductor quantum dots (SQDs), are notable for showing significantly high fluorescence; however, their toxic nature (due to heavy metals’ presence), they are not appropriate for biological applications such as bioimaging, biosensing and drug delivery. In contrast, carbon dots (CDs) have zero-dimensional structure, excellent fluorescent properties and nearly negligible toxicity, which gives them an advantage over other fluorescent carbon-based nanomaterials.^3^

Many different types of carbon sources have already been used for the synthesis of CDs. Both bottom-up and top-down approaches have been explored. In the top-down approach, the large carbon materials are cut down to small fine nanoparticles using laser ablation or acid etching.^4^ The bottom-up approach takes place by fusing the organic molecules by either carbonization or dehydration using the hydrothermal/solvothermal/thermal synthesis method.^5^ Among all the methods hydrothermal method proved to be the best method for simplistic and fast synthesizing CDs. The as-prepared CDs synthesized shows fluorescence in the yellow region. L-ascorbic acid, urea and ortho-phenylenediamine (o-PDA) were used as precursor materials for the low-cost synthesis of CDs using the hydrothermal method. To achieve fluorescence in the visible region, we use o-PDA; as o-PDA is an organic molecule, the CDs synthesized may not be very stable in water. L-ascorbic acid is used to attach more functional groups on the surface of CDs so as to make them more stable in water. For nitrogen doping of CDs, both urea and o-PDA are used as both these compounds have nitrogen groups present in their molecular structure. Handful papers are being published on CDs showing fluorescence in the visible region. Sathish and co-workers have synthesized yellow CDs using O-PDA and pyrazole using the carbonization method. The yellow fluorescent CDs synthesized show maximum emission at 415 nm excitation wavelength and have a quantum yield of 10.3% used for heavy metal sensing in water samples.^6^ Xiaojuan et al. have synthesized yellow fluorescent phosphorus and nitrogen co-doped CDs from pumpkin peel and phosphoric acid (H_3_PO_4_) with satisfactory quantum yield and used them for bioimaging purposes.^7^ Although very few yellow fluorescent CDs have been reported, the synthesis method is tedious or the product obtained is in small amount, limiting the use of CDs for bioimaging purposes. The morphology of nanoparticles also plays a part in their biomedical applications.

Herein, the yellow fluorescent CDs we have synthesized through the one-step hydrothermal method are cost-effective and straightforward with a yield of 40% and a fluorescent quantum yield of 8 %. To use the CDs for biological applications, *in vitro* cytotoxicity tests were performed to understand whether these nanoparticles are compatible for cellular imaging as well as *in vivo* model systems like zebrafish.^8,9^ Zebrafish is rapidly gaining interest in biomedical applications because of its high degree of similarity with human genome and fast generation time.^10,11^ The high fecundity and optical transparency of the zebrafish larvae allow real-time studies of the bioimaging test and give them an advantage over other animal models such as mice and rats.^12–15^ The uptake studies were done in 72 hours post fertilized (hpf) zebrafish larva. We also performed cellular studies to check through which pathway these CDs enter inside the mammalian cells and for bioimaging applications, these studies are essential to understand how the CDs interact with the cells and how they are uptaken by the cells and 3D animal model. The yellow fluorescent CDs enter the cells through a particular endocytic pathway called Clathrin-mediated endocytosis (CME). Overall, the CDs we have synthesized show promising results and can be used as a good bioimaging tool in the future.

**Scheme 1:**
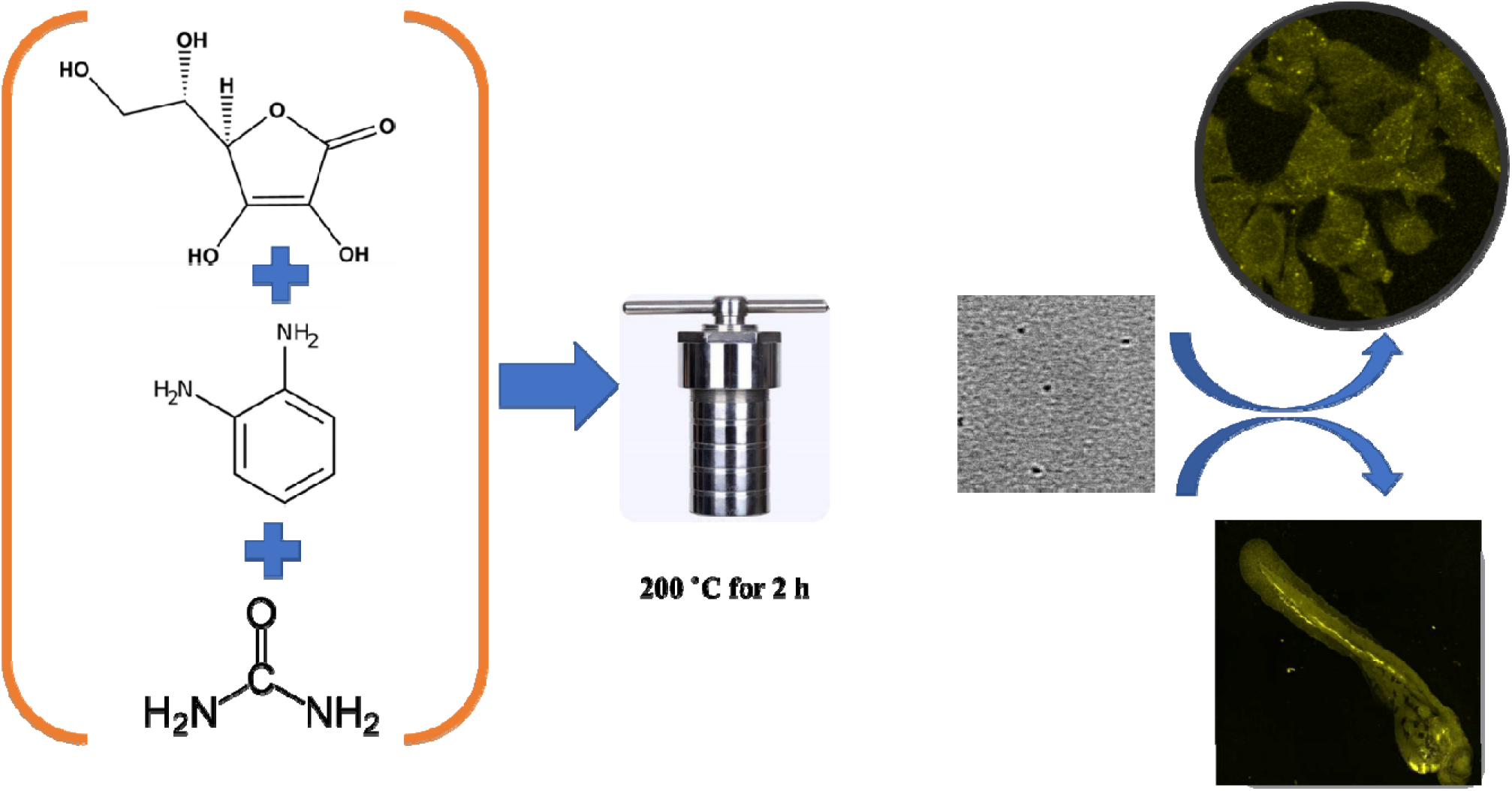
CDs synthesis schematic representation. Ortho-phenylenediamine, l-ascorbic acid and urea dissolved in Milli-Q water are used for synthesis, autoclaved at 200□C for 2 h leading to the one-step formation of yellow fluorescence emitting carbon dots that could be used for bioimaging application by uptake of these nanoparticles in mammalian cells and zebrafish larva.

## 2. Results and discussion

### 2.1 Synthesis and characterization of CDs

Carbon dots showing yellow fluorescence are synthesized using the hydrothermal synthesis method, and a simple bottom-up approach is used to synthesize CDs. In detail, 0.2 g l-ascorbic acid, 0.05 g ortho-phenylenediamine (OPDA), and 0.05 g urea were mixed in 14 ml of deionized water, placed in a Teflon container, and heated in a microwave oven at 200°C for 2h. After cooling the hydrothermal setup, the dark brown solution obtained is filtered through a syringe filter (0.22 micron) to remove all the larger particles. The pH of the CD solution is maintained at 7.4 (physiological pH) using sodium hydroxide (NaOH) and then lyophilized to obtain powder CDs (brown color) to be further used for different characterization and experiments. Transmission electron microscopy (TEM) and atomic force microscopy (AFM) were used to understand the morphology and structure of CDs. The TEM image of CDs shown in figure 1a shows the average size of CDs is 2.7 ± 1.5 nm **(figure 1a,b)**. A total of 100 CDs’ diameters were calculated through image J software to get an approximate value of the size of CDs. The Atomic Force microscopic (AFM) characterization shows the topological height of CDs comes out to be 0.88 nm (**figure 1c,d)**. The characterization of CD’s structural composition was done using X-ray diffraction (XRD) and Fourier transform infrared spectroscopy (FTIR, **figure 1e,f)**. We obtain a few sharp peaks in XRD spectra, proving that CDs are polycrystalline.^16^ We get very different peaks for the precursor materials as compared to CDs in the XRD spectra, proving that the CDs formed to have other structural properties as compared to the precursor materials used **(figure 1e)**. The FTIR spectra give a qualitative idea of the types of functional groups attached to the nanoparticle’s surface. The main elements present are hydrogen (H), carbon (C), nitrogen (N) and oxygen (O), forming chemical bonds (-OH) stretch, (-NH) stretch and (-C=C) aromatic stretch at 3326 cm^-1^, 2929.57 cm^-1^ and 1625.19 cm^-1^ as shown in **figure 1f**. Also, the amino group presence confirms by obtaining a peak at 2029.57 cm^-1^, which also proves that CDs are N-doped. The presence of stretch (-C=C) at 1625.19 cm^-1^ shows that CDs contain an aromatic group of carbon, and this may be due to the use of ortho-phenylenediamine (OPDA) as precursor material.

**Figure 1.**
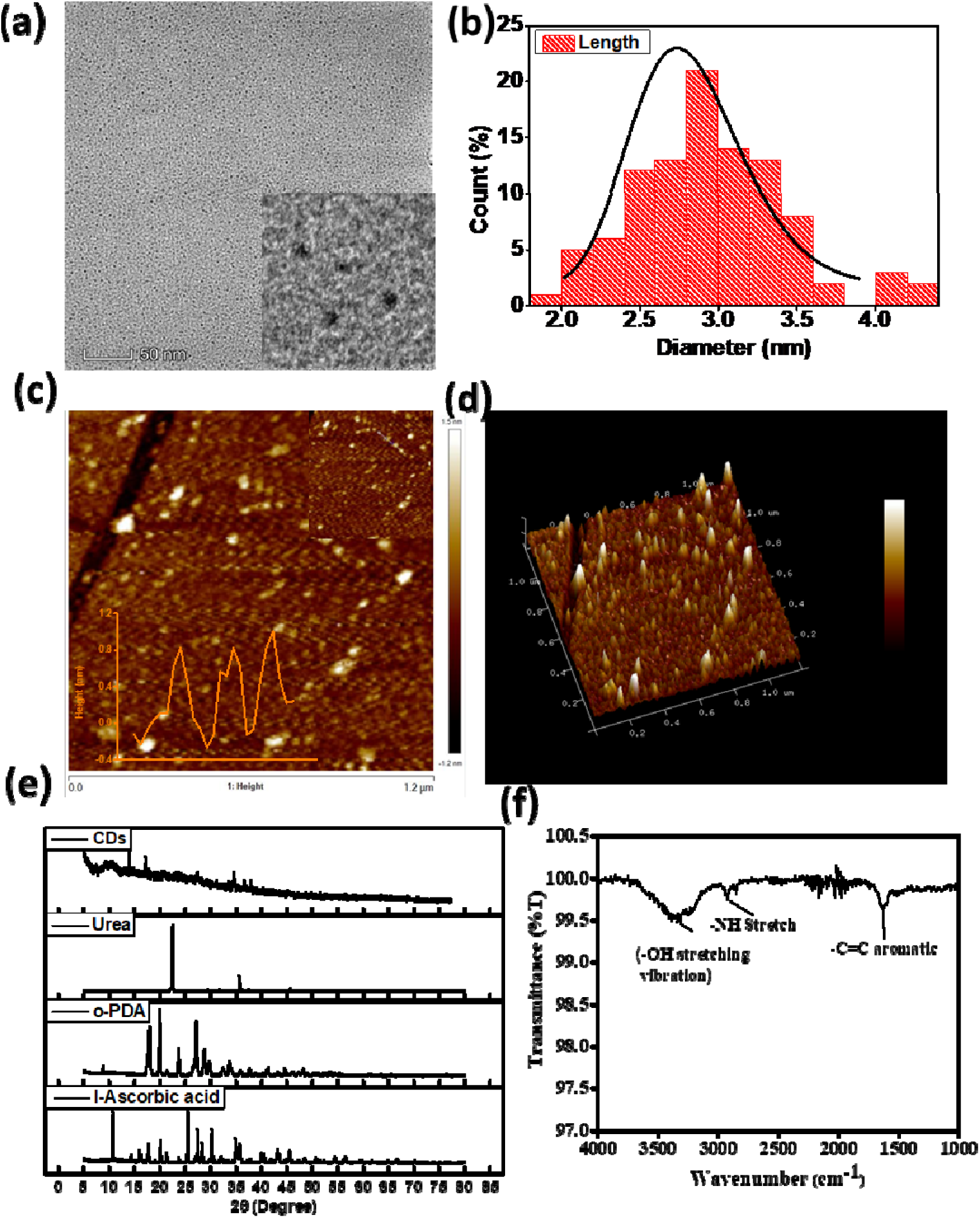
Characterization of CDs **(a)** Transmission electron microscopy (TEM) image shows spherical-disk shaped CDs of size 2.7 ± 1.5 nm (inset showing zoomed images of CDs) **(b)** Statistical size distribution of CDs calculated through image J software **(c)** Atomic force microscopy (AFM) image of CDs showing the disc-shaped morphology of CDs with the topological heights of 0.88 nm which is approx. equal to 3-4 layers of graphene sheet **(d)** 3D image of CDs taken using AFM **(e)** X-ray diffraction (XRD spectra of the CDs and the precursor materials used for the synthesis. The sharp peaks show that CDs are crystalline in structure **(f)** The functional groups attached to the surface of CDs are confirmed by Fourier transform infrared (FTIR).

### 2.2 Optical properties of CDs

CDs’ optical properties are studied using UV-visible absorption spectra and fluorescence spectra obtained by UV-Visible spectrometer and fluorescence spectrophotometer. In UV-visible spectra, we get three absorbance peaks 240, 280 and 320 nm. The absorbance peak attained at 240 nm is designated to the aromatic carbon structure’s π-π* transition. At 280 nm the peak obtained is assigned to the π-n* transition. The transition happened from functional groups having lone pairs of electrons. As the lone pair of electrons does not directly affect the π-π* transition, they are directly involved in the π bonding of the aromatic system. There is a relevant intramolecular charge transfer for the planer amines and aromatic rings in the CDs leading to broader fluorescence spectra.^17^ The absorbance peak attained at around 320 nm wavelength due to energy level transition from the conjugated π-structure of angstrom size present in CDs **(figure 2a)**.

**Figure 2:**
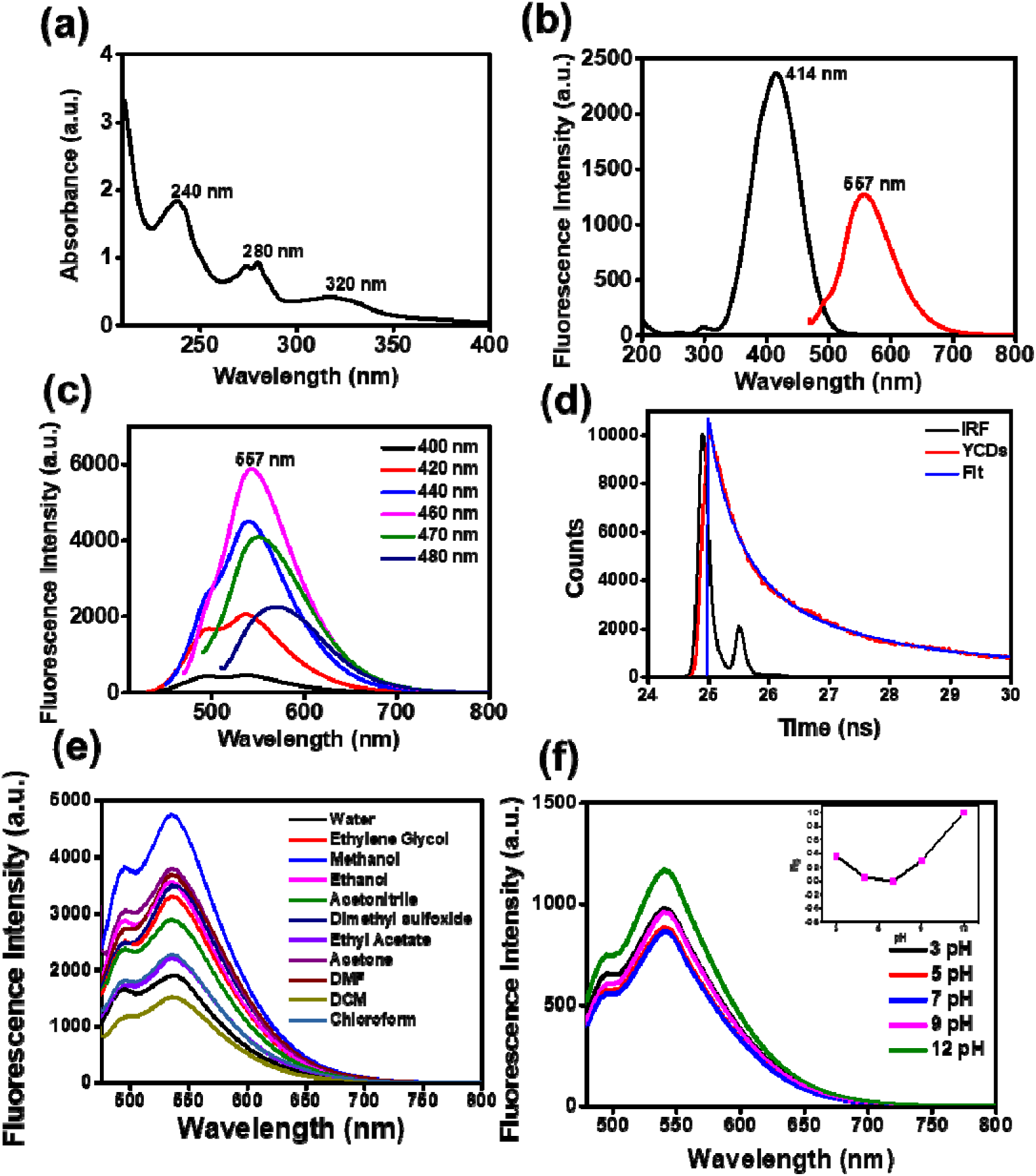
(a) Ultraviolet (UV)-Visible spectra of CDs showing three absorbance peak-240 nm, 280 nm and 320 nm because of different levels of transition (b) Graph showing the excitation and emission spectra of CDs at 460 nm excitation wavelength showing highest emission peak at 542 nm wavelength (c) CDs excited at different excitation wavelength ranging from 400 nm - 480 nm with the highest emission obtained at 460 nm excitation wavelength (d) Lifetime plot of yellow fluorescent CDs, calculated through time-correlated single-photon counting (TCSPC) technique. The average lifetime of CDs comes out to be (1.69 ± 0.95 ns). (e) CDs dispersed in different solvents (Water, Ethylene glycol, Methanol, Ethanol, Acetonitrile, Dimethyl sulfoxide (DMSO), Ethyl acetate, Acetone, Dimethylformamide (DMF), Dichloromethane (DCM), Chloroform) showing different emission intensities at a concentration of 0.5mg/ml (f) CDs’ emission spectra dispersed in different pH solutions (3,5,7,9 and 12). The fluorescence intensity is stable until pH nine but increases at pH 12.

Fluorescence spectra of yellow CDs show a maximum emission peak at 557 nm with an excitation wavelength of 460nm. There is no shift in the emission peak with the change in excitation wavelength, as shown in **figure 2b**. The CDs were excited in the wavelength range of 400 nm to 480 nm **(figure 2c)**, specifying that CDs synthesized have independent fluorescence behavior that is not observed often. The CDs show yellow color in white light and no fluorescence in UV light when dispersed in an aqueous solution.

The quantum yield of the fluorescent nanoparticle gives the fraction of excited molecules that have returned to the ground state with the release of emission photons; in other words, the quantum yield is the ratio of photons emitted to the number of absorbed photons.^18^ The fluorescence emission studies were executed using the FP-8300 Jasco spectrofluorometer (Japan) in the excitation range of 400 to 480 nm. 10 mm path length quartz cuvettes were used with a slit width of 10 nm to record the emission spectra of CDs. The reference standard for the calculation of quantum yield of CDs was taken fluorescein (L=0.92 in 0.1 M NaOH solution) based on the excitation spectra of CDs. The CDs’ fluorescent quantum yield comes out to 8% in milli-Q water by referring to standard fluorescein. The synthesized CDs are more dispersed in protic solvents such as alcohols because of forming a hydrogen bond between the nitrogen atom and solvent molecule. This leads to an inversion of the lowest-lying n-π* state to π-π*, which leads to higher fluorescence quantum yield.^17^ The fluorescence lifetime measurement was taken at room temperature using a picosecond time-correlated single-photon counting (TCSPC) setup (Edinburgh instrument Ltd, Life spec II model). A picosecond light-emitting diode laser is used at 515 nm for the excitation of CDs. The average lifetime of the CDs is 1.69 ± 0.95 ns **(figure 2d)**. Both quantum yield and lifetime are characteristics of significant importance in the case of fluorescent nanoparticles used for biomedical applications. The larger the fluorescence quantum yield easier it is to detect the CDs used as a fluorescent probe. Many parameters can affect quantum yield and fluorescence lifetime, such as temperature, pH, polarity, viscosity, hydrogen bonding, presence of quencher etc. All these parameters should be checked before using CDs as the fluorescent probe.

The powdered CDs obtained after lyophilization were again dispersed in different solvents-water, ethylene glycol, acetone, methanol, ethanol, acetonitrile, dimethyl sulfoxide (DMSO), ethyl acetate, acetone, dimethyl formamide (DMF), dichloromethane (DCM) and Chloroform, arranged in descending order based on their polarity. The concentration of CDs was kept fixed at 0.5mg/ml. The solvents are organized based on the decrease in polarity of the solvent. As the CDs are dispersed in different solvents, their emission spectra do not shift with a change in solvent polarity. Still, there is an increase in fluorescence intensity when CDs are dispersed in more non-polar solvents than water. The highest emission intensity is obtained when methanol is used as a solvent. This may be because protic solvents such as alcohol make a hydrogen bond with the amine group present on the surface of the CDs leading to an inversion of the lowest-lying n-π* state to π-π*, which gets down to higher fluorescence quantum yield **(figure 2e)**.

For stability of these nanoparticles in various biological milieu, pH-dependent studies of CDs were performed. The CDs were placed in different pH solutions, and changes in their fluorescence signal as a function of pH were recorded. The pH solutions of 3,5,7 & 12 pH were prepared in Milli-Q water. CDs (0.5 mg/mL) were added to these different pH solutions, and emission spectra were taken, as shown in **figure 2f**. The fluorescent CDs offer an equilibrium fluorescence value till pH 9, but after that, very high-intensity fluorescence was found at pH 12. The reason behind the pH sensitivity of CDs can be explained by the protonation and deprotonation of oxygen-containing groups on the surface.^19^ There is no notable colorimetric change of CDs dispersed in different pH solutions ranging from pH 3 to 12, indicating their robust optical stability.

The use of CDs for the bioimaging applications of their fluorescence stability in various concentrations of an ionic solution is significant. CDs’ ionic stability was checked in different potassium chloride (KCl) solution concentrations are prepared (0.1-1M). Our results show that CDs’ fluorescence is relatively stable in different concentrations of KCl. There is a slight decrease in fluorescence intensity of CDs, but it’s not significant **figure 3a**. CDs’ photostability was also checked by treating the CDs solution in milli-Q water with continuous excitation wavelength for 30 mins, and readings were recorded after every 5 min. The results show an insignificant decrease in fluorescence with time, designating good photostability of CDs as shown in **figure 3b**. A thermal stability study of CDs was also performed using a spectrophotometer by taking fluorescence reading at different temperatures (10°C to 80°C). There is an insignificant decrease in fluorescence intensity of CDs with an increase in temperature, as shown in **figure 3c**. The CDs were also kept at 80°C for 30 mins; the decrease in fluorescence intensity is not very significant as stipulated through the (I/Io) graph **(figure 3d)**. All these studies show that the CDs synthesized are pretty stable. We also studied the metal ion stability of these fluorescent CDs. The CDs shows significant fluorescence stability in the presence of heavy metal ions, as shown in the **supplementary figure S2**. Hence, CDs are a good candidate for bioimaging as they are not sensitive to heavy metal ions present in traces inside the body, which are essential for human metabolism.

**Figure 3:**
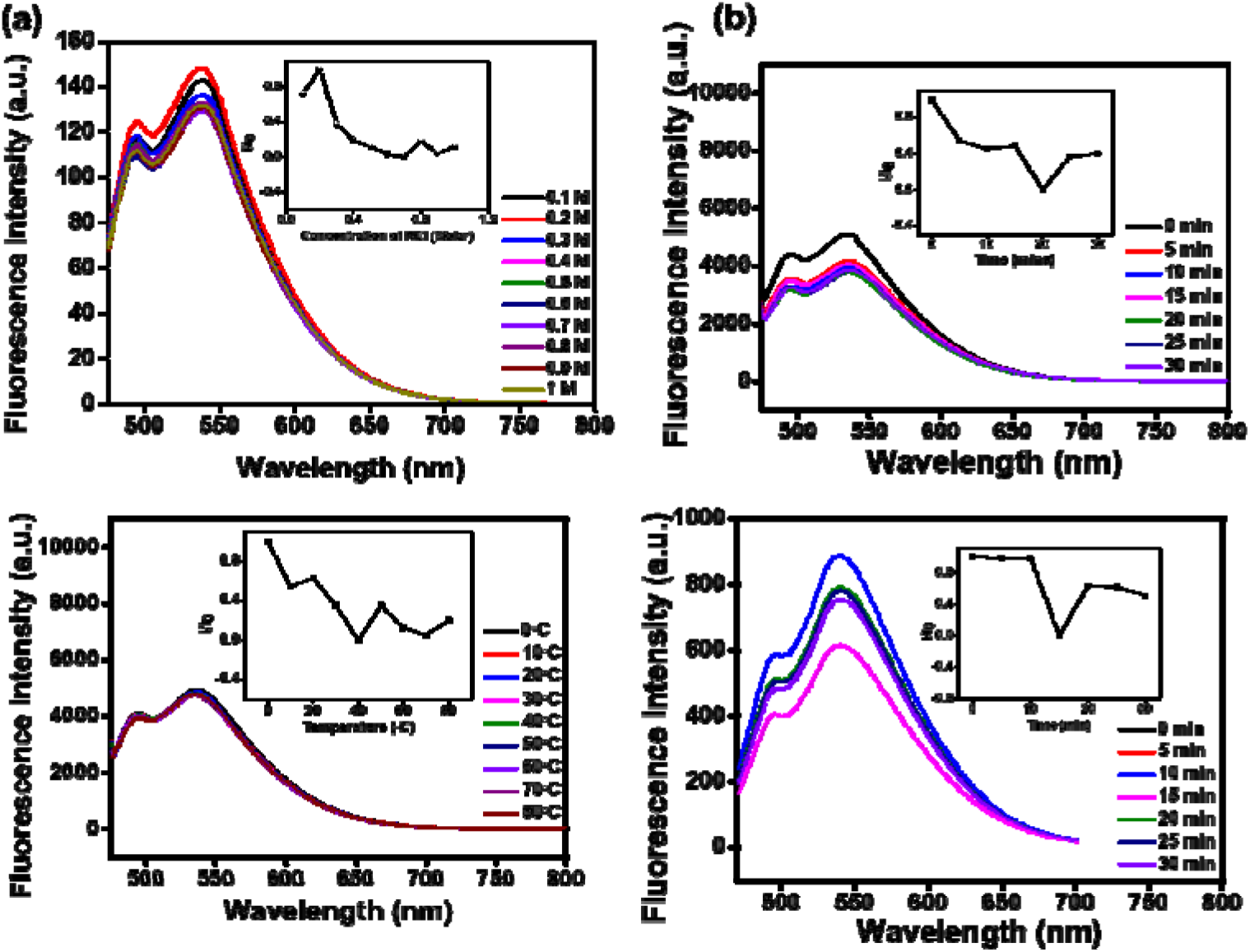
Fluorescence stability of CDs **(a)** CDs’ emission spectra recorded after treated with different ionic concentrations of potassium chloride (KCL) (0.1 to 1M) **(b)** CDs’ emission spectra recorded at every five minutes in over 30 min timepoint, treated with an excitation wavelength of 460 nm continuously to check their photostability. **(c)** CDs’ fluorescence spectra taken at different temperature ranges (10°C-80°C), at an increment of 10°C to check the effect of temperature of fluorescence intensity. **(d)** CDs’ fluorescence spectra recorded at different time points (0-30 min) after keeping the solution at 80 □C. The relative effect on the ionic stability, photostability, and thermal stability of CDs excited at 460 nm is shown in the inset.

### 2.3 Cellular uptake studies of CDs

To develop biocompatible, non-toxic CDs for their application in bioimaging, we performed a (3-(4, 5-dimethyl thiazolyl-2)-2, 5-diphenyltetrazolium bromide) (MTT test) checking the viability of the cells in the presence of CDs. We have selected two cell lines, one is the human embryonic kidney (HEK), and another one is the retinal pigment epithelial (RPE) cell line. Approximately 2 × 10^4^ cells were seeded in each well of a 96-well plate and grew to confluent stage. To measure the output from both the cell-line assays we use a luminescence-based microplate reader (SpectraMax 190 Microplate Reader, Molecular Devices). Cells which were no exposed to CDs were kept as negative control in both imaging test and other biological assays. In the MTT experiments different concentrations, namely 20, 40, 60, 80, 100, 120, and 150 µg/mL of CDs were used. In RPE cells, after 24 h of incubation with CDs, the experiment showed ∼100% cell viability at a concentration 60 µg/mL of CDs and then decreased to ∼70% up to 150 µg/ml concentration of CDs (Supplementary Figure **S1**). In the case of the HEK cell line, MTT for 1h period, almost all the cells were viable, but for the 24hr period, the cell viability starts decreasing with an increase in QD concentration. The cytocompatibility studies on RPE and HEK cells suggested that CDs show different levels of toxicity in different cell lines. As the CDs show the best biocompatibility in RPE cells, we continued cellular uptake studies at different concentrations. Cells were incubated with CDs of varying concentrations for 30 mins at 37°C, and then the cells were washed and fixed. Confocal microscopy-based uptake studies of the samples show successful internalization of the CDs in RPE cells. Through confocal images, we have quantified the data suggesting that the CDs exhibit strong intracellular fluorescence as the concentration increases. It is proved through concentration-dependent studies that with increase in concentration of CDs from 50 to 150 µg/ml the fluorescence intensity also increases **figure 4a,b**. We checked for their fluorescence in other channels and found negligible emission in other channels except in the yellow channel.

**Figure 4:**
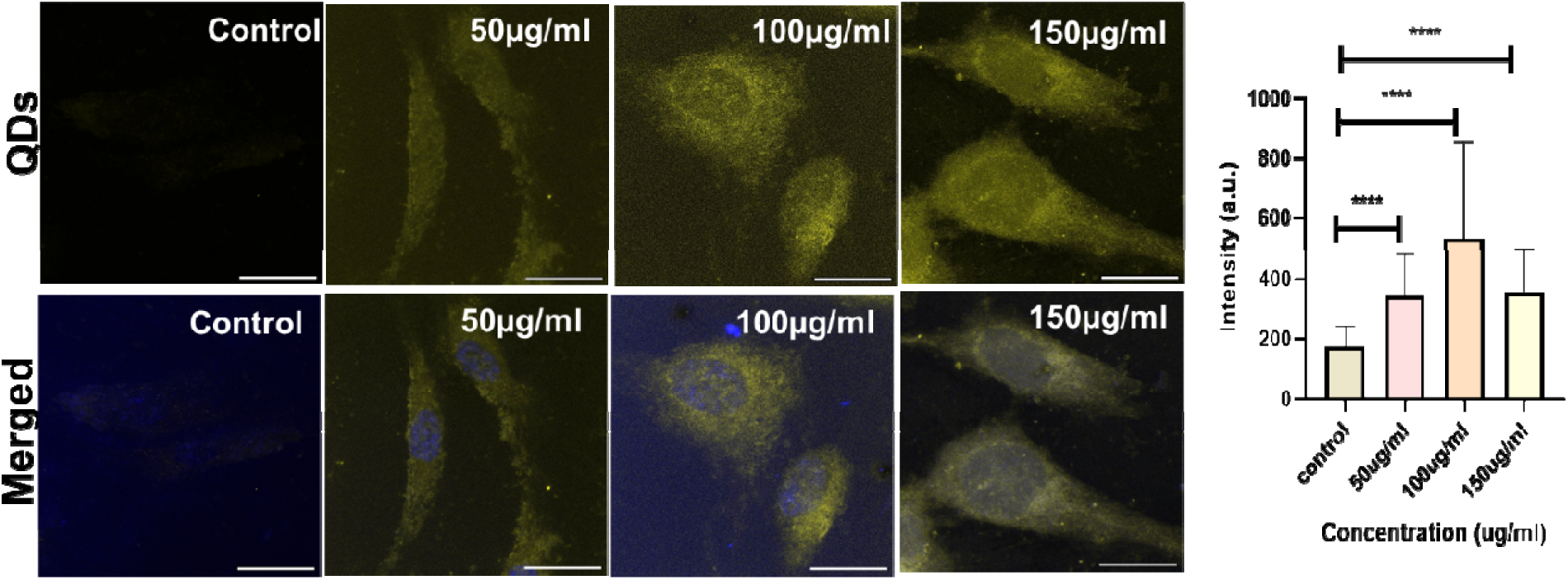
Cellular uptake studies of CDs based on increase in concentration of CDs, using Confocal images of RPE cells incubated with 50, 100 and 150 µg/mL of CDs, respectively. The RPE cells were imaged after treating the cells with CDs for 30 min. The scale bar is 20 µm for all the images. Cellular uptake of CDs at 50,100 and 150µg/mL was also quantified using Fiji ImageJ software .**** Indicates statistically significant value of p < 0.0001, ** indicates statistically significant value of p = 0.002. and * = 0.01 (t-test).

To understand the endocytic pathway through which the CDs were internalized inside the cells, we performed an endocytic pathway response experiment. We use the SUM-159 breast cancer cell line for the experiment. Two important pathways for cellular internalization of nanoparticles are Clathrin-mediated endocytosis (CME) and clathrin-independent endocytosis (CIE). We have chosen two cellular cargoes for the studies; Transferrin (Tf), a notable marker for CME, and Galectin3 (Gal3) for labelling CIE. Transferrin receptor (TR) is a crucial iron transporter controlling iron homeostasis as well as an important marker for CME.^20^ Gal3 triggers glycosphingolipid (GSL)-dependent biogenesis of different classes of endocytic structures called clathrin-independent carriers (CLICs) hence follow CIE pathway.^21^ Both these proteins were used as a reference control for the experiment. Specific inhibitors were used to inhibit the two main pathways. Pitstop is used to inhibit CME, and lactose is used to inhibit the CIE pathway. Both these inhibitors were used to understand the uptake of CDs. The cells were treated with these marker proteins and CDs in the presence of inhibitors. We observed that CDs and transferrin uptake are significantly reduced by pitstop treatment (**figure 5a,b)**. At the same time, lactose treatment does not significantly affect the fluorescence of transferrin and CDs, whereas Gal3 uptake was reduced considerably. We have also checked the CIE pathway uptake of CDs using Gal3 as a reference marker. Extracellular Gal3 is inhibited by lactose which competes for the same β-galactosides on the plasma membrane receptors, inhibiting the uptake of extracellular Gal3.^13^ It was observed that on treatment with lactose, Gal3 uptake in cells was reduced significantly, whereas CDs uptake decreased is not very significant. Pitstop, when used as an inhibitor, the uptake of Gal3 inside the SUM-159 cells decreased marginally, but the uptake of CDs decreased significantly (**Figure 6a,b,c)**. These results confirm that CDs uptake is through the CME pathway.

**Figure 5:**
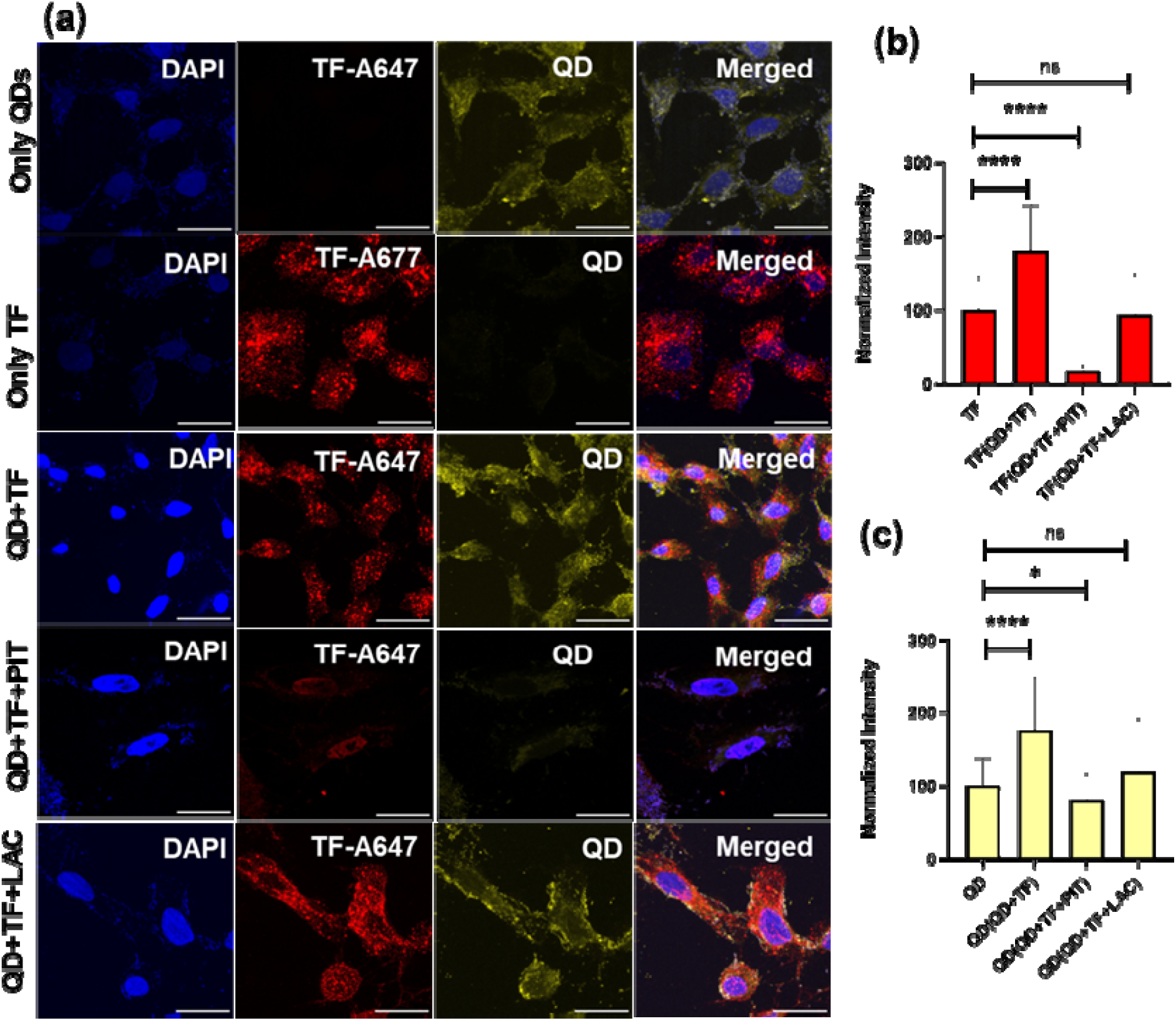
Uptake of CDs in SUM-159 cells via clathrin-mediated endocytosis. (a) Uptake of A647 labelled transferrin (Tf) in SUM-159 cells in the presence of CDs when exposed to pitstop-2 (20µM) and lactose (100mM) (b) Showing the quantification analysis of normalized intensity of Transferrin (Tf). (d) Showing the quantification analysis of normalized intensity of CDs. The scale bar is calibrated at 20 µm for all the images. **** denotes the statistically significant p-value (p□0.0001), whereas ns indicates a statistically non-significant p-value. (One–way ordinary ANOVA). A total of 30 cells and n = 2 independent experiments were performed.

**Figure 6:**
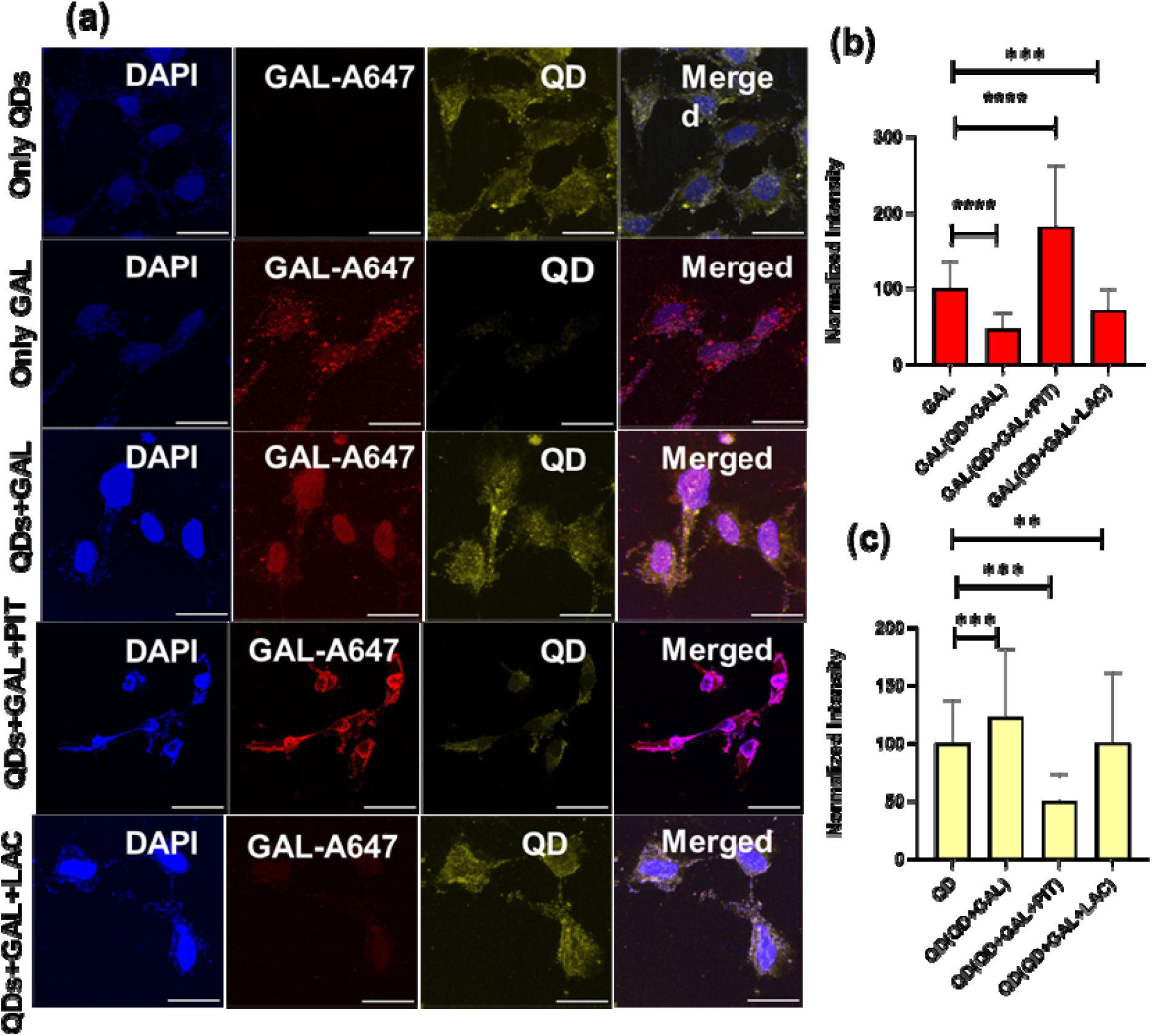
Uptake of CDs in SUM-159 cell line via clathrin-mediated endocytosis. (a) Uptake of A647 labelled Galectin (GAL) in SUM-159 cells in the presence of CDs when exposed to pitstop-2 (20µM) and lactose (100mM) (b) Showing the quantification analysis of normalized intensity of Galectin (GAL). (d) Showing the quantification analysis of normalized intensity of CDs. The scale bar is calibrated at 20 µm for all the images. **** denotes the statistically significant p-value (p□0.0001), whereas ns indicates a statistically non-significant p-value. (One–way ordinary ANOVA). A total of 30 cells and n = 2 independent experiments were performed.

### 2.4. CDs uptake in zebrafish model system

After performing the concentration-dependent cellular studies and understanding the mechanism of cell internalization, we have studied the *in vivo* imaging potential of CDs using zebrafish larva as an animal model. Scientists have already shown that the CDs enter the zebrafish larvae body through skin absorption and accumulate, mainly in the yolk sac.^22,23^ The zebrafish chosen for the experiment were Assam wild type. Salinity, pH and oxygen level are appropriately maintained to get healthy fertilized eggs. After the eggs hatching, the larva was collected for 72 hpf for the experimental study. The CDs were exposed to zebrafish larva for different time points (2, 4 and 6 hrs.) and at different concentrations (200, 300 and 400 µg/ml) **(supplementary information figure S3)**. The uptake of CDs increases with an increased time. The uptake of the CDs is highest in the yolk sac and the tail region of the larvae. The highest uptake of CDs is observed at 4 hr. time point, and then it decreases at 6 hr. time point. The zebrafish studies show that the CDs are easily uptaken by zebrafish without showing any toxicity. The CDs offer bioaccumulation properties and further confirm these CDs’ bioimaging potential. In **figure 7a,b,c,d,e** we see at 200 ug/mL fixed concentration the fluorescence intensity increases with increase in time point. Through the zoomed image of zebrafish larvae **(figure 7f)** we can configure that there is high fluorescence emission at the yolk sac region and notochord of a zebrafish larva.

**Figure 7.**
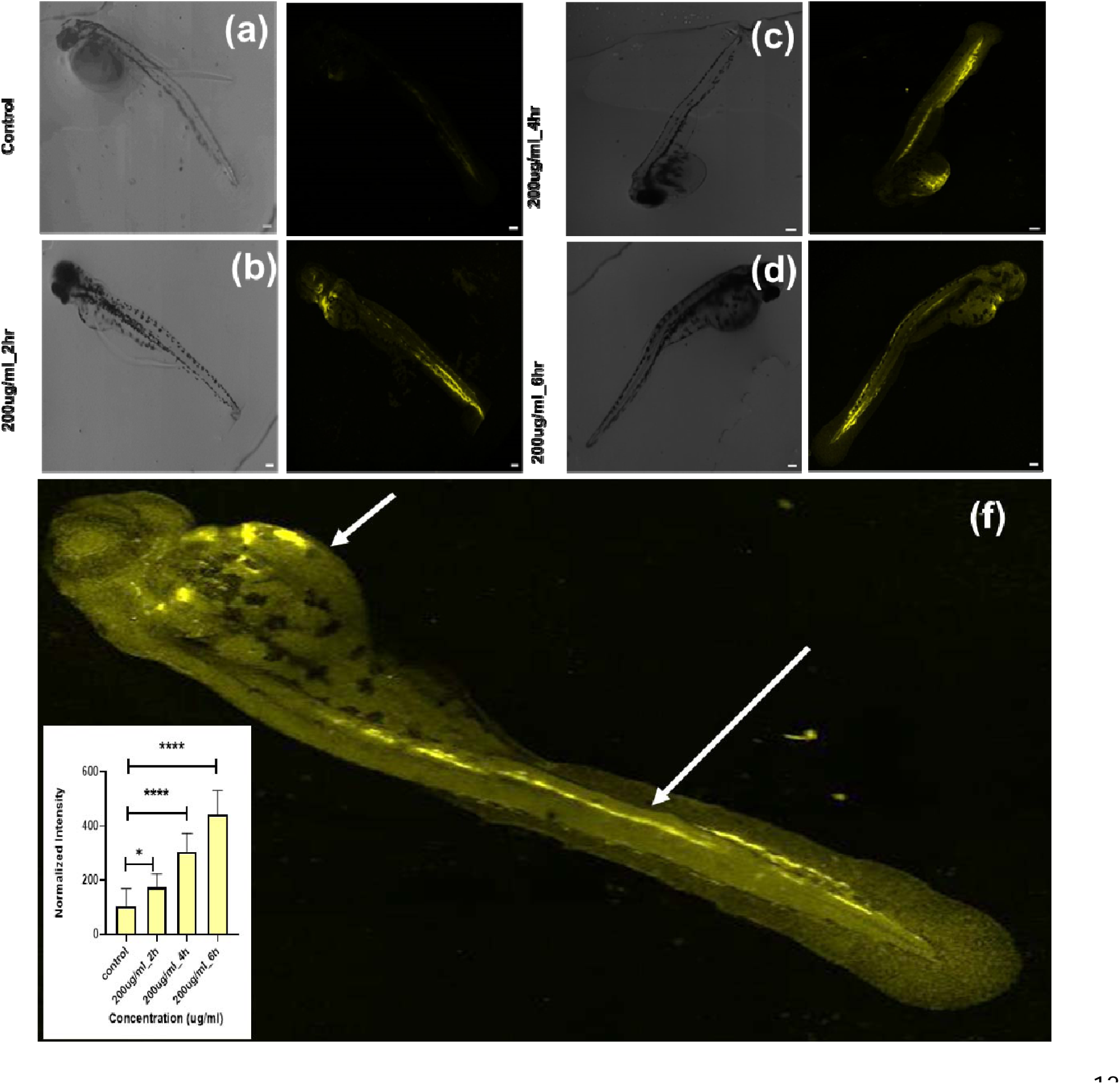
Uptake study of CDs in zebrafish larva. 200µg/ml of CDs uptaken by 72 hpf zebrafish larva and the treatment performed for different time points (2, 4 and 6 h) and also at different concentration (200, 300 & 400 µg/ml) (f) (shown in inset) Quantification analysis of the fluorescence intensity of CDs uptaken by 72hpf zebrafish larva for different time period (concentration of CDs are fixed at 200µg/ml). The scale bar calibrated at 100µM. **** denotes the statistically significant p-value. (p < 0.0001). *Denotes the statistically significant p-value (p < 0.05). 08 larva per condition were quantified. (f) a representative zoomed-in image of a zebrafish with tissue-specific uptake of CDs.

### 2.5 Scratch test to study cell proliferation

Many ligands and nanoparticles trigger signaling events at the plasma membrane during endocytosis which triggers different physiological processes like cell migration. To check the effect of uptake of CDs on collective cell migration, we performed a scratch test assay to study the wound healing properties of CDs. Retinal pigment epithelial (RPE) cells were used for the wound healing study. This cell line has been already used for cell migration studies.^25^ RPE cells were grown in 6 well plates, and at 100% confluency, a scratch is made using 200µL tip which mimics an actual wound. The treatment of cells with CDs of different concentrations was experimented, and images were taken at different time points to analyze the cells’ migration. The cells migrate properly even when the CDs were present. At the concentration of 20 μg/mL and 50 μg/mL concentration of CDs, the cell migration is higher than in control, proving that CDs are biocompatible and help wounds heal faster, as shown in **figure 8 (a,b)**.

**Figure 8.**
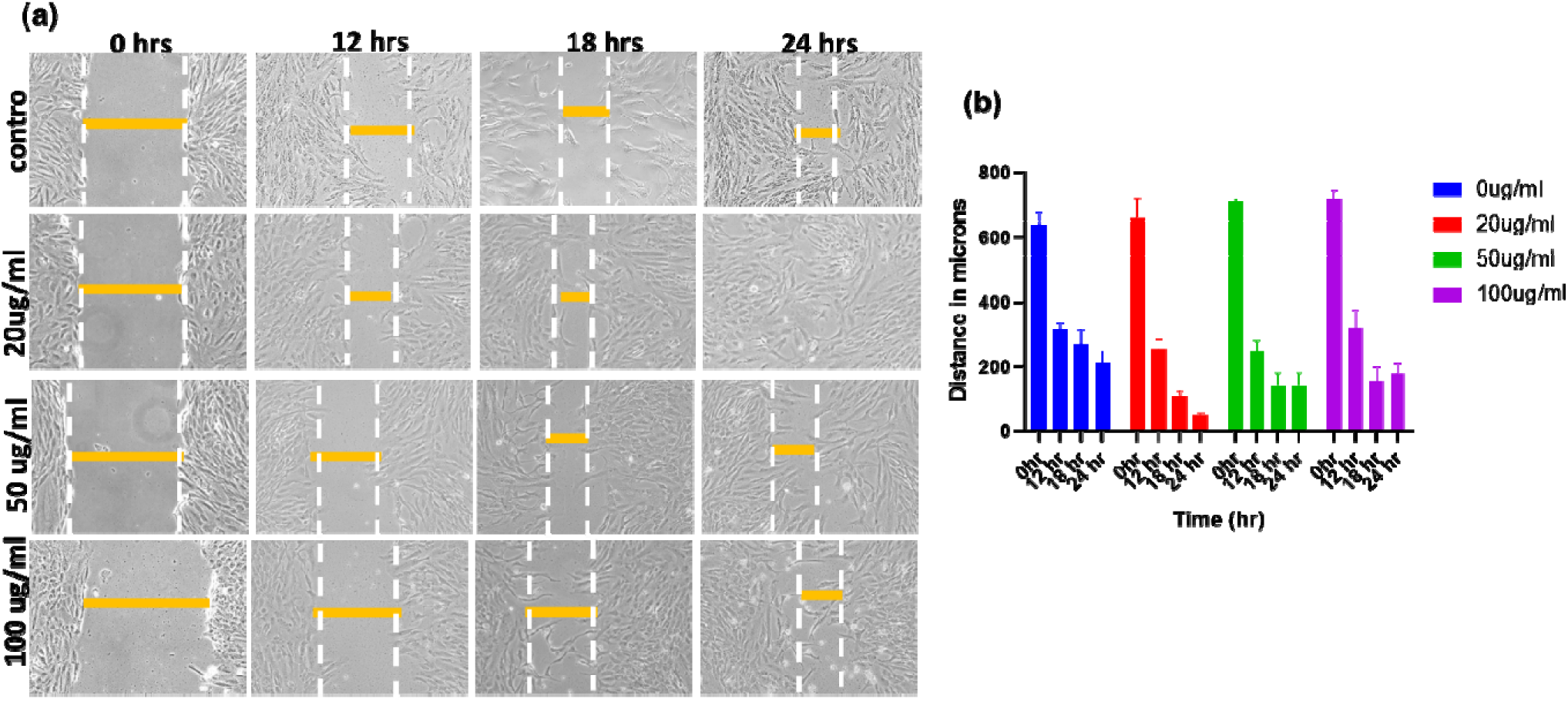
**(a)** Images taken using Nikon camera showing wound healing process using RPE cells in control, 20, 50 and 100 μg/mL of CDs at different time points. (b) Graph showing how the induced wound heals faster in the 20 and 50 μg/mL concentration compared to control. The wound healing is almost the same in control and at 100 μg/mL of CDs.

## 3. Conclusions and discussions

We synthesized yellow emitting carbon dots (CDs) through the hydrothermal method using ortho-phenylenediamine (OPDA) and urea as precursor material which also helps in the nitrogen doping of CDs. L-ascorbic acid is also used to react with carbon colloids to produce oxygen and nitrogen-containing surface defects on CDs and is associated with fluorescence emission of the particle.^26^ Milli-Q water is used as the solvent. The synthesized CDs are stable in a water medium and can be easily used for biological applications. The CDs synthesized were characterized through different techniques analyzing the structural and optical properties of CDs. With the excitation wavelength of the light region of the electromagnetic spectrum, there is negligible harm to the cells when yellow fluorescence emitting CDs are excited as compared to most of the CDs excited in the ultraviolet(UV) region, killing the cells by rupturing cell’s DNA. The quantum yield of CDs comes out to be 8% in water. The lifetime of QDs comes out to be 1.69 ± 0.95 ns. The fluorescence of QDs also depends on the solvent they are dispersed in, and highest fluorescence we obtain in methanol. The peak doesn’t shift, but the fluorescence intensity increases from Milli-Q water to methanol. The fluorescence intensity is also dependent on pH. The fluorescence intensity is almost stable till 9 pH and intensity increases drastically at pH 12. The CDs are stable in an ionic solution, photostability and thermal stability were also checked, and the fluorescence CDs are remarkably stable in all the conditions. Even in the presence of different heavy metal ions, the fluorescence intensity of CDs didn’t quench. The cytotoxicity of CDs was also checked using RPE and HEK cell lines. The RPE cells were almost 70-80% viable in a 100ug/ml concentration of CDs. Both in vitro and in vivo bioimaging experiments were done by treating cells and zebrafish larvae with different concentrations of CDs and at different time points. We also find out that CDs were uptaken through clathrin-mediated endocytosis inside the cells. The current work provides a promising fluorescent nanomaterial for bioimaging and can also be used by modulating the surface properties and conjugating it to different biomolecules, and using them for other biomedical applications of CDs in the coming times.

## 4. Materials & methods

### 4.1 Materials

Ortho-phenylenediamine (OPDA), urea, l-ascorbic acid, mowiol, Hoechst, pitstop(PIT), Galectin3 (Gal3) (Alexa 647), Transferrin (Tf) -A647, 3-(4,5-Dimethylthiazol-2-yl)-2,5-diphenyltetrazolium bromide (MTT), were obtained from Sigma-Aldrich. Acetone (>99.5%), N,N-dimethylformamide (>99.5%), ethanol(>99.5%), methanol (>99.8%), acetone, acetonitrile, Isopropyl alcohol(IPA), Ethyl acetate, Dimethylformamide (DMF), Dichloromethane (DCM), Chloroform), syringe filter and lactose were acquired from Merck. Dimethyl sulfoxide (DMSO), fluorescein, Paraformaldehyde, and cell culture dishes for adherent mammalian cell culture were obtained from Himedia. Dulbecco’s modified Eagle’s medium (DMEM), Ham’s Nutrient Mixture F12 (HAMs F12), fetal bovine serum (FBS), trypsin-EDTA (0.25%) and penicillin-streptomycin from Gibco. All the chemicals were of good scientific quality, and no further sterilization or treatment is needed.

### 4.2 Synthesis of Fluorescent CDs

Yellow fluorescent emitting CDs were synthesized by the reflux method. Ortho-phenylenediamine (OPDA) and urea are used as precursor material dissolved in 14 ml of Milli-Q water. L-ascorbic acid is used as a stabilizing agent for CDs as it promotes the formation of defects on the surface of CDs and the formation of functional groups on the surface, enhancing CDs’ fluorescence intensity. The total time of reaction is two hours at 200 °C. After providing heat as a source of energy for the formation of CDs, the autoclave reactor is permitted to cool down to room temperature. The CDs synthesized were separated from the bigger particles through a syringe filter (0.02 mm). The original CD solution has a slightly acidic pH which brings it back to physiological pH using a 0.1 M sodium hydroxide (NaOH) solution. The CDs containing the solution are lyophilized, and we obtain the brown powder, which is dissolved in an appropriate medium for further experimental use.

### 4.3 Analytical methods used to study the characterization of CDs

To analyze the shape and size of fluorescent CDs FEI Titan Themis transmission electron microscope (TEM) is used. The atomic force microscope (Multimode 8 (Bruker)) took both two-dimensional and three-dimensional images of CDs. X-ray diffraction studies were done to characterize the crystalline nature of the CDs. The scans were recorded with a speed of 0.2°/min from 5° to 80° with Cu-K_α_ radiation in Bruker-D8 DISCOVER X-ray spectrometer. FP-8300 Jasco spectrofluorometer (Japan) is used to excite the CDs in the 400 to 500 nm range. Fluorescein was used as a standard reference for relative quantum yield measurement of CDs, which comes out to be 8% in water. UV-VIS absorbance spectra of CDs were acquired by UV-VIS spectrophotometer Spectro Cord-210 Plus Analytic Jena (Germany). In ATR mode, the Fourier transform infrared spectroscopy (FT-IR) of CDs was recorded using the FTIR spectrometer from Spectrum 2, PerkinElmer. Scanning was done in the range of 1000 cm^-1^ to 4000 cm^-1^. The bioimaging of fixed cell samples and the fixed zebrafish larvae was done under 63X and 10X resolution using confocal laser scanning microscopy platform Leica TCS SP8.

### 4.4 Spectroscopic studies

The recording of both absorbance and emission spectra was done using spectroscopic grade solvents. The recording of the emission spectra takes place 10nm after the excitation wavelength for the standard recording of emission spectra. Fluorescein in 0.1 M NaOH solution (□ = 0.92) is used as a reference for calculating the relative quantum yield of CDs. The following equation for quantum yield calculation:

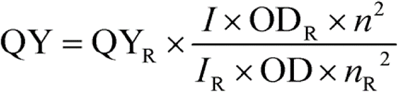

Where QY is the quantum yield, *I* is the integrated fluorescence intensity, OD is the optical density, *n* is the refractive index, and R denotes for reference.

### 4.5 Cell Culture

Mesenchymal triple-negative breast cancer cell line (SUM159A), Retinal pigment epithelial (RPE) cells and Human embryonic kidney (HEK) cells were gifted by Prof. Ludger Johannes Institut Curie, Paris, France. SUM159A cells were cultured in HAMs F12 complete media. RPE and HEK cell lines were cultured in DMEM complete media. PBS of 1X strength with pH 7.4 was used for all the studies.

#### 4.5.1 MTT assay

The cytotoxicity of the CDs was analyzed so as to use them for bioimaging applications. 96-well plates were used to seed the cells. The cell density was maintained at approx. 8,000 cells per well (100 µl) and kept in the incubator for 24 h in a CO_2_ incubator maintained at 5% CO_2_ and 95% humidity for proper cell growth. The incubated cells after 4 h were washed with PBS before treatment with CDs.100 µl of 7 different concentrations of CDs viz., 20, 40, 60, 80, 100, 120 and 150µg/mL triplicates and were incubated for 1h and 24 h. 3- (4,5-dimethylthiazol-2-yl)-2,5-diphenyltetrazolium bromide (MTT) colorimetric assay was used after incubation of cells, to quantify the effect of CDs on cell viability. The cells were washed twice with PBS after the treatment with CDs; then, 10 μL of MTT solution (5 mg/mL) was added to each well, and the solutions were further incubated for 3 h at 37°C. The formazan crystals formed after the incubation were further dissolved in 100 μL DMSO and kept in the dark for 15 min by wrapping the plate with aluminum foil. The intensity of the color was then measured by the absorbance at 570 nm from the formazan crystals and was representative of the number of viable cells per well. The values thus obtained for the untreated control samples were equated to 100%, and the relative percentage values for CDs were calculated accordingly. All experiments were performed in triplicate, and the cell viability (%) was calculated using the following equation:

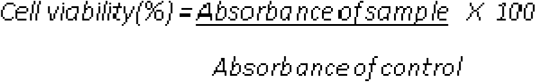

#### 4.5.2 Confocal Microscopy Studies

Two different cell lines - SUM 159 and RPE were grown in HAMS-F12 and DMEM cell culture media at a density of ∼40,000 cells/well on coverslips placed in 24 well plate cell culture plates. The cells were kept in incubator for 24 h at 37 °C and 5% CO_2_. After getting approximately full confluency, cells were washed twice with PBS and treated with different concentration of CDs (50, 100 and 150 µg/mL). The treated cells were further incubated for 30 min, respectively. After 30 mins of incubation the cells were washed with PBS twice before fixing them with 4% PFA for 20 mins. After treatment of cells with 4% PFA for 20 mins, cells were washed with PBS thrice to remove all the left-over particles and proper removal of 4%PFA. The fixed cells were then stained with Hoechst, and mounted on glass slide using a mounting medium (mowiol). Finally, the Leica TCS SP5 confocal microscope using 63X oil immersion objective were used to take images to study the cellular internalization of CDs. The excitation source of Hoechst is 4o5nm laser, while a 488 nm argon laser was used as an excitation source for CDs. The emission bandwidth for Hoechst and CDs was 410-450 nm and 500-630 nm, respectively.

#### 4.5.3 Cellular uptake studies via clathrin mediated endocytosis

Cellular studies were performed to investigate the pathway by which CDs were entering inside cells. Briefly, 24 well plate was used to seed SUM-159 cells on coverslips with cell density of approximately 40,000 cells/well. The cells were treated with pitstop (20 μM) and lactose (100 mM) in HAMs-F12 (serum-free media) in seeded wells and incubated for a particular time period at 37 °C. Two main pathways studied till date for cell endocytosis clathrin-mediated endocytosis (CME) and clathrin-independent endocytosis (CIE) pathways. Pitstop used to inhibit CME pathway and lactose is used to inhibit CIE pathway. Non-treated cells were used as control. After the treatment the cells were washed carefully with PBS and treated with fluorescently labelled markers, which followed CME (Transferrin-A647(5 μg/mL)) and CIE (Galectin-A647 (5 μg/mL)) pathways and CDs (100 µg/mL) for a specific time interval at 37°C. The cell fixation was done using 4 % PFA at 37°C for 15 mins and washed three times with PBS to remove any left PFA or any kind of impurity. The coverslips taken from each well were mounted properly on mowiol containing Hoechst for imaging. The image of the fixed cells was taken using a confocal laser scanning microscope under 63X resolution. Fiji ImageJ software was used for the quantification study of the images obtained using a confocal microscope.

#### 4.5.4 *In vivo* studies in Zebrafish larvae

Based on the organization for economic cooperation and development (OECD) guidelines, the uptake studies were performed on 72hpf zebrafish larva. The 72 hours post fertilized live larvae were placed in six-well plates, and dead larvae were removed. In one well, 15 larvae were placed. Each set of the larva is treated with CDs at different time points (2h, 4h and 6h) and at different concentrations (200, 300 and 400 µg/mL) **(supplementary figure S3)**. One well was set as a control where larvae were not treated with nanoparticles. E3 media is used as a medium after CD treatment and is used to wash the larva twice to remove the excess CDs. The larvae were fixed using 4% PFA solution (used as a fixative solution for 2-3 mins). Post-fixation, the larvae were mounted on a glass slide using the mounting solution Mowiol. The zebrafish larva slides were dried first and then used for further confocal imaging analysis.

## Acknowledgements

We sincerely thank all the members of the DB group for critically reading the manuscript and for their valuable feedback. US thanks IITGN-MHRD, GoI for PhD and Dr Ankit Gangrade for helping in cell culture experiments. KS thanks GSBTM for the postdoctoral fellowship. KK thanks SERB, GoI, for National Postdoctoral Fellowship. DB thanks SERB, GoI for Ramanujan Fellowship, IITGN for the startup grant, and DBT-EMR, Gujcost-DST & GSBTM for research grants. Imaging facilities of CIF at IIT Gandhinagar are acknowledged.

## Supplementary Information

**Figure SI 1:**
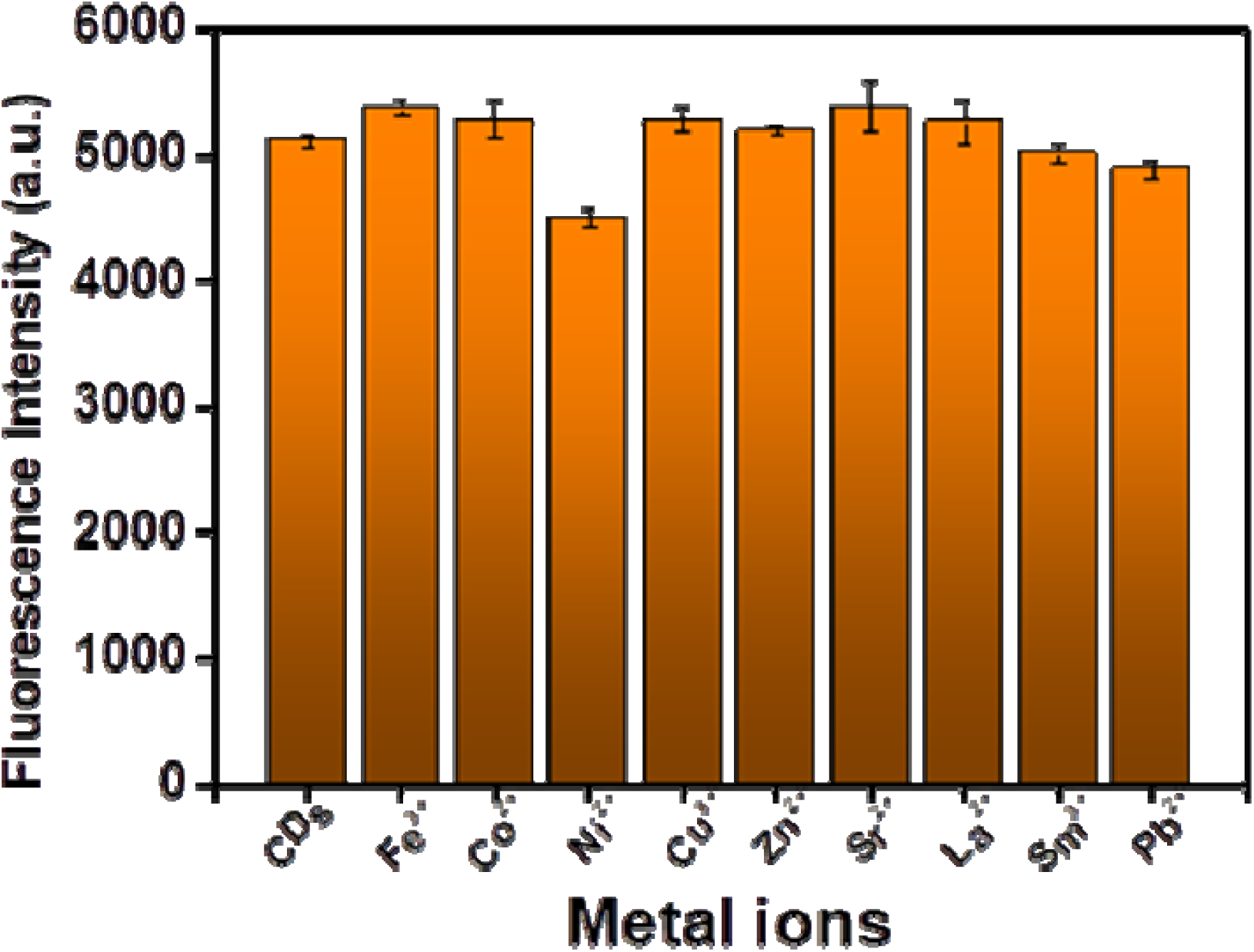
Fluorescence intensity of CDs in the presence of different heavy metal ions. The fluorescence intensity of CDs does not affect significantly in the presence of different metal ions.

**SI 2:**
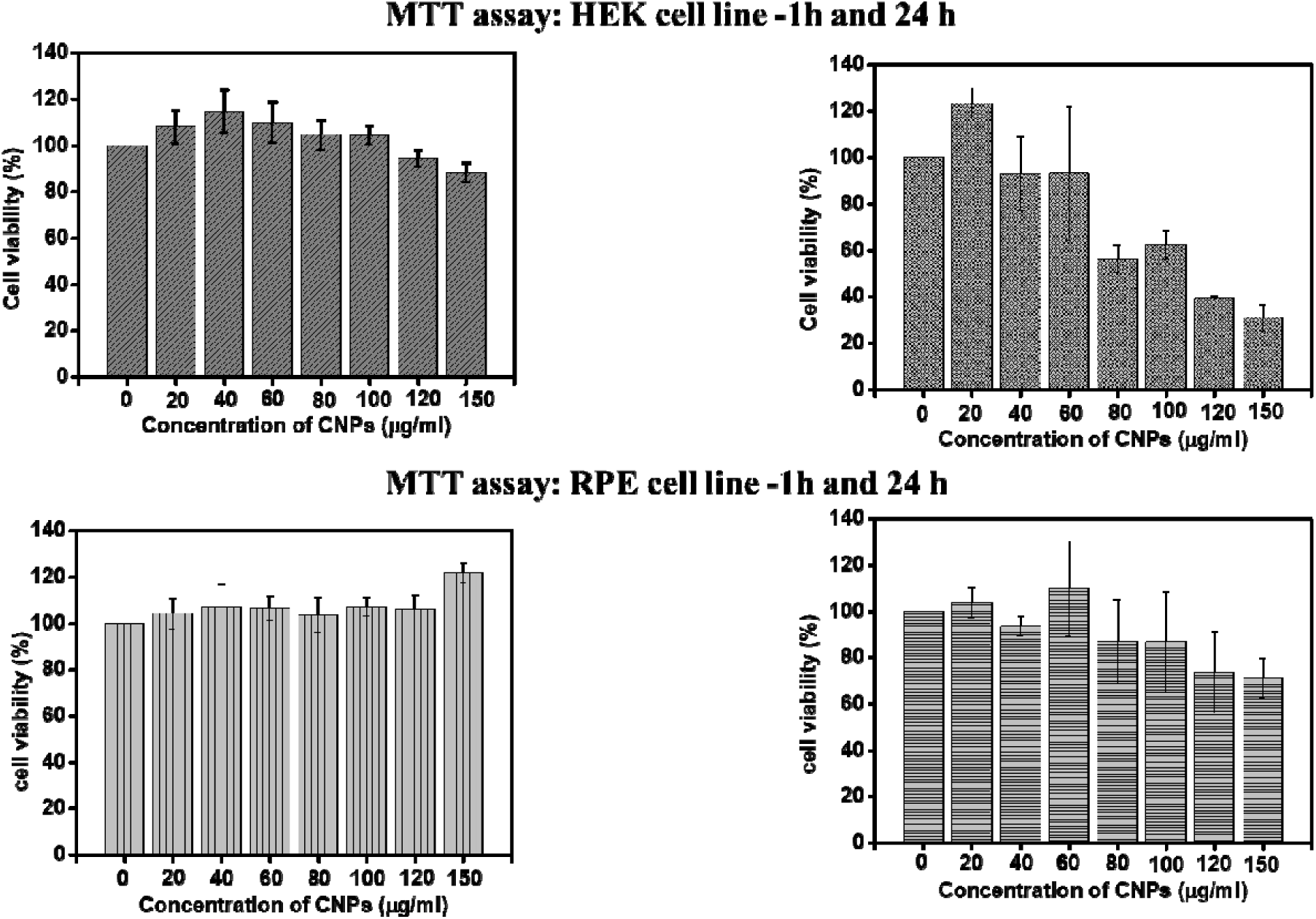
MTT assay of CDs in two different cell lines-Human embryonic kidney (HEK) and Retinal pigment epithelium (RPE) cell line.

**Figure S3:**
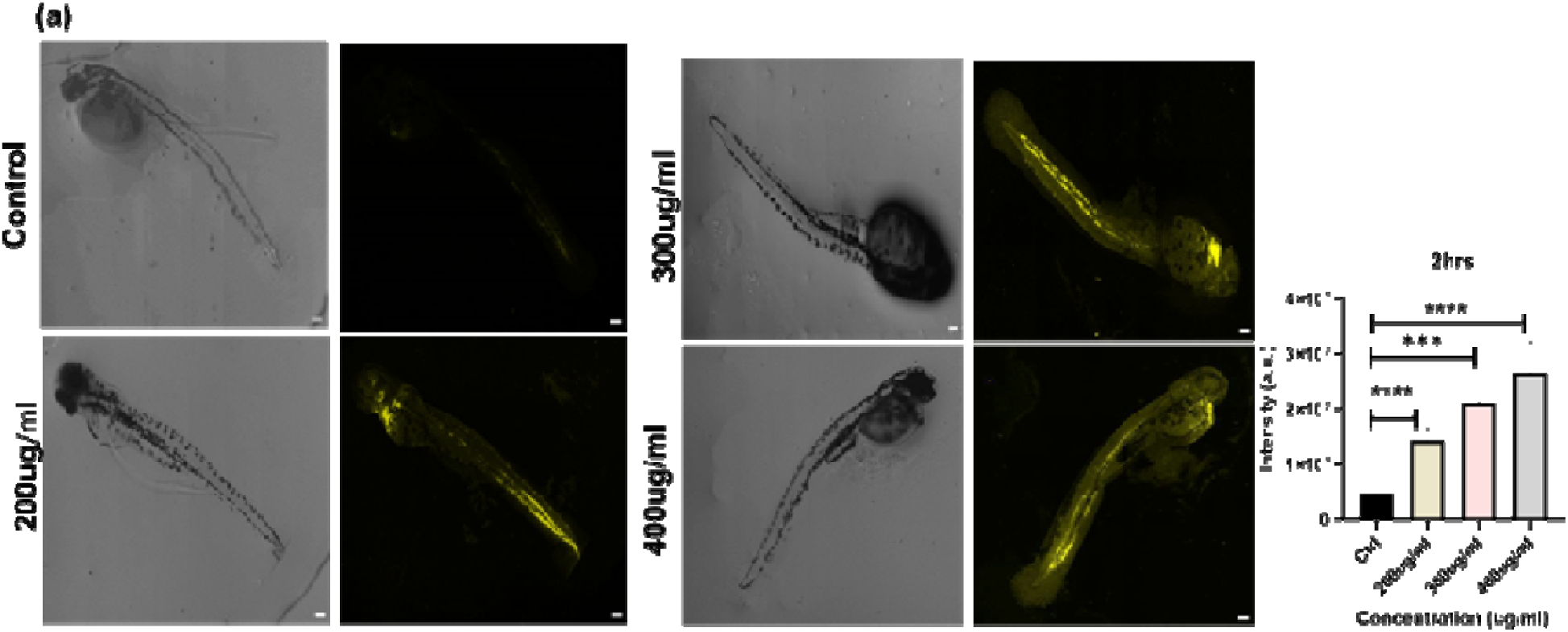
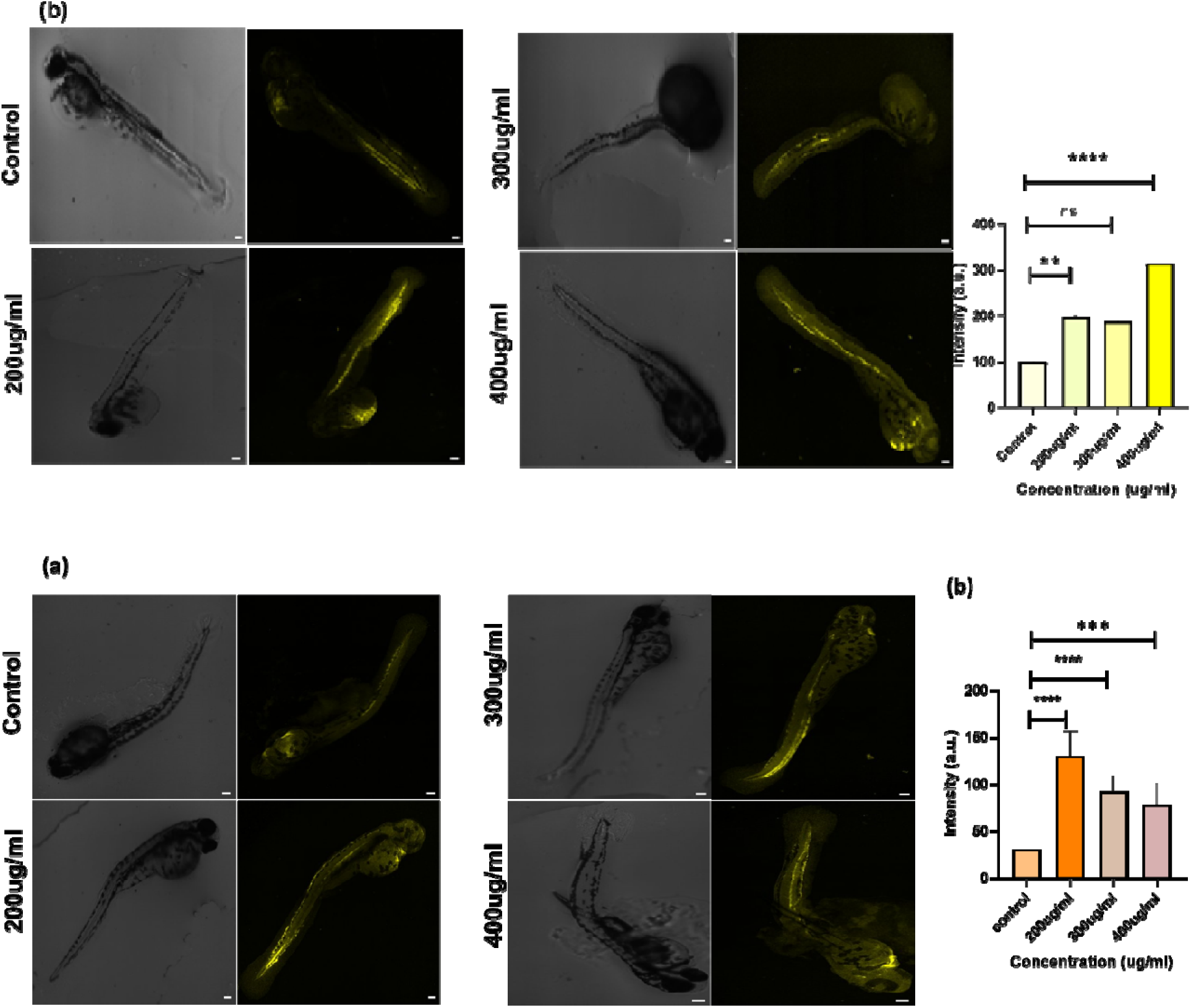
(a-c) Confocal images of zebrafish larva at different concentrations (200, 300 and 400 µg/mL) at different time point (2, 4 and 6h); graphs are showing fluorescence intensity of CDs uptaken by zebrafish larvae for different concentration of CDs at fixed time point. The scale bar is set at 100µM. **** denotes the statistically significant p-value. (p < 0.0001). *Denotes the statistically significant p-value (p < 0.05). 08 larva per condition were quantified.

